# Direct profiling of genome-wide dCas9 and Cas9 specificity using ssDNA mapping (CasKAS)

**DOI:** 10.1101/2021.04.16.440202

**Authors:** Georgi K. Marinov, Samuel H. Kim, S. Tansu Bagdatli, Alexandro E. Trevino, Josh Tycko, Tong Wu, Lacramioara Bintu, Michael C. Bassik, Chuan He, Anshul Kundaje, William J. Greenleaf

## Abstract

Detecting and mitigating off-target activity is critical to the practical application of CRISPR-mediated genome and epigenome editing. While numerous methods have been developed to map Cas9 binding specificity genome-wide, they are generally time-consuming and/or expensive, and not applicable to catalytically dead CRISPR enzymes. We have developed a rapid, inexpensive, and facile assay for identifying off-target CRISPR enzyme binding and cleavage by chemically mapping the unwound single-stranded DNA structures formed upon binding of a sgRNA-loaded Cas9 protein (“CasKAS”). We demonstrate this method in both *in vitro* and *in vivo* contexts.

---

CRISPR-based methods for editing the genome and epigenome have emerged as a highly versatile means of manipulating the genetic makeup and regulatory states of cells. CRISPR technologies hold the potential to transform medical practice by enabling direct elimination of pathogenic sequence variants or manipulation of aberrant gene expression programs. CRISPR has also become a standard tool for discovery in biomedical research, including its uses for high-throughput, massively parallel genomic screens ^1^.

The presence of significant off-target effects is of universal concern for genome engineering technologies, presenting a major hurdle to fully realizing their potential utility. CRISPR tools have been shown to exhibit biochemical activity away from their intended target sites, which is particularly problematic for therapeutic applications, where risks of activity at sites other than the intended target leading to negative consequences to patient health must be minimal. Understanding and mapping these effects is therefore an urgent need.

To this end, numerous experimental approaches have been developed to experimentally map off-target effects genome-wide. Methods such as Digenome-seq ^2^ look for particular types of cut sites around target sequences in whole-genome sequencing data; however, deep whole-genome sequencing remains expensive. Assays such as BLESS ^3^, GUIDE-seq ^4^, HTGTS ^5^, DSBCapture ^6^, BLISS ^7^, SITE-seq ^8^, CIRCLE–seq ^9^, TTISS ^10^, INDUCE-seq ^11^, and CHANGE-seq ^12^ aim instead to directly map Cas9 cleavage events. However, all these methods involve some combination of complex and laborious molecular biology protocols and non-standard reagents, and have not been widely adopted. Other methods, such as DISCOVER-seq ^13^, which maps DNA repair activity by applying ChIP-seq against the MRE11 protein, as well as earlier applications of ChIP-seq to map catalytically dead dCas9 occupancy sites genome-wide ^14,15^, suffer from technical issues associated with the ChIP procedure. Most recently, long-read sequencing has been adapted to the problem of Cas9 specificity profiling, in the form of SMRT-OTS and Nano-OTS ^16^, but the cost of these methods is relatively high while their throughput is comparatively low.

Various computational models have also been trained to predict off-targets genome-wide ^17,18^. However, these exhibit far from perfect accuracy, and thus in many situations, especially within clinical contexts, direct experimental evidence is needed to accurately identify potential unintended effects of CRISPR-based reagents.

A faster, more accessible, and versatile method for mapping CRISPR off targets is thus still a critical need in the field. When a Cas9-sgRNA ribonucleoprotein (RNP) is engaged with its target site, the sgRNA invades the DNA double helix, forming a ssDNA structure on the other strand (Fig. 1a). We thus reasoned that mapping ssDNA-containing regions should be a sensitive biochemical signal of productive Cas9 binding. The recently developed KAS-seq ^19^ assay for mapping single-stranded DNA (ss-DNA) (**k**ethoxal-**a**ssisted **s**sDNA sequencing ^19^) is ideally suited for the purpose of identifying ssDNA generated by CRISPR protein binding to DNA (Fig. 1a-b). KAS-seq is based on the specific covalent labeling of unpaired guanine bases with N_3_-kethoxal, generating an adduct to which biotin can then be added using click chemistry. After shearing, biotinylated DNA, corresponding to regions containing ssDNA structure, can be specifically enriched for and sequenced.

**Figure 1:**
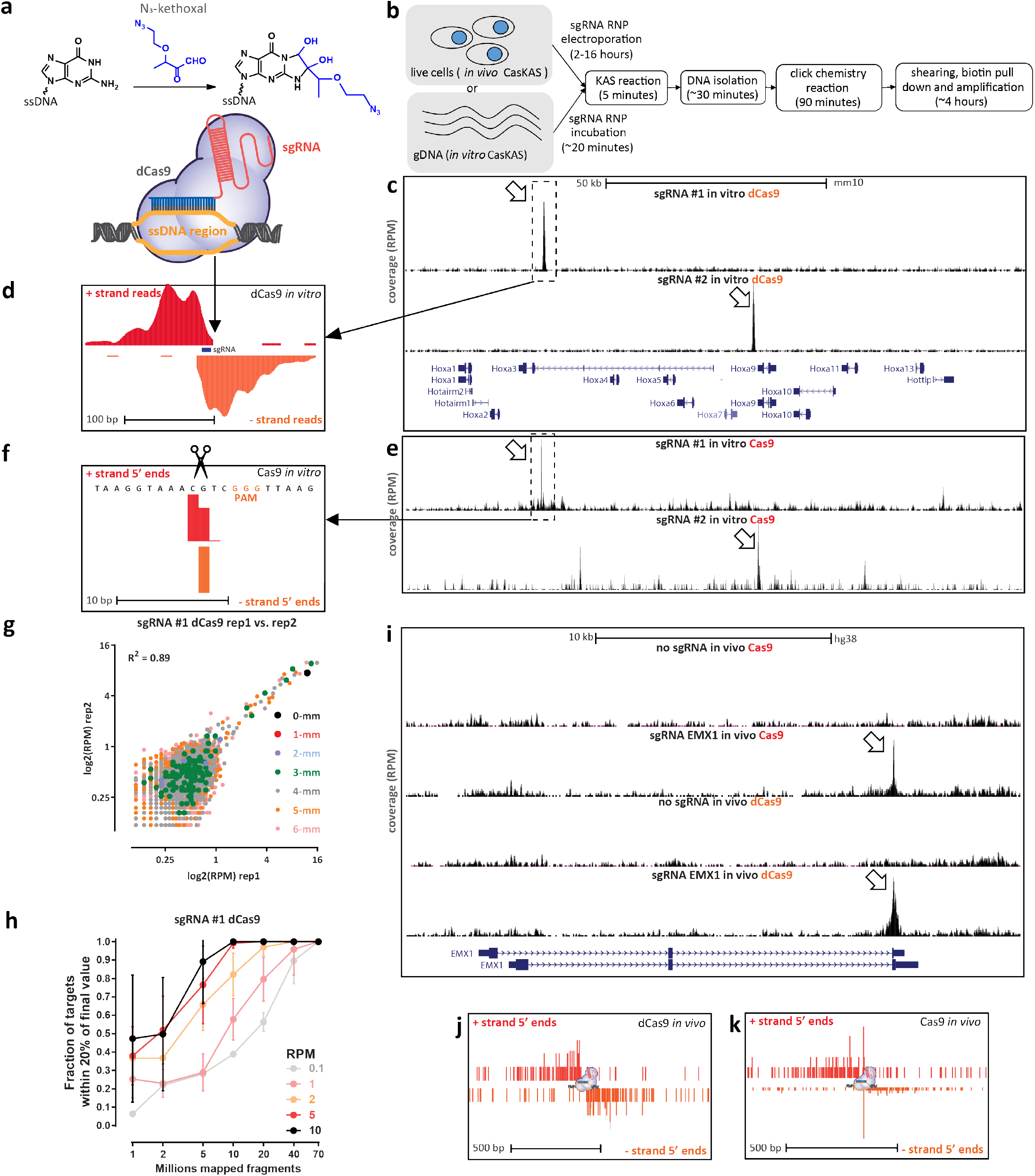
CasKAS maps dCas9- and Cas9-mediated strand invasion and cleavage events genome-wide *in vitro* and *in vivo*. (a) CasKAS is based on the KAS-seq assay for mapping ssDNA structures. N_3_-kethoxal covalently modifies unpaired guanine bases (while having no activity for G bases paired within dsDNA). Strand invasion by Cas9/dCas9 carrying an sgRNA results in the formation of a ssDNA structure, which can be directly identified using N_3_-kethoxal. (b) Outline of *in vivo* and *in vitro* CasKAS. For in *in vitro* CasKAS, gDNA is incubated with a dCas9/Cas9 RNP, then N_3_-kethoxal is added to the reaction; for in *in vivo* CasKAS, cells are transfected with an RNP, then treated with kethoxal. DNA is then purified, click chemistry is carried out, DNA is sheared, labeled fragments are pulled down with streptavidin beads, and sequenced. (c and d) Mapping of dCas9 targets *in vitro*. (c) Mouse gDNA was incubated with dCas9 RNPs carrying one of two sgRNAs targeting the mouse *HOXA* locus. Highly specific labeling is observed at the expected target location of each sgRNA. (d) Asymmetric strand distribution of *in vitro* dCas9 CasKAS reads around the sgRNA target site. (e and f) Mapping of Cas9 targets *in vitro*. (e) Mouse gDNA was incubated with Cas9 RNPs carrying one of same two sgRNAs targeting the mouse *HOXA* locus. (f) The distribution of 5’ read ends around targets sites in *in vitro* CasKAS datasets shows direct capture of the intermediate cleavage state. (g) Reproducibility of *in vivo* dCas9 CasKAS datasets. Shown are RPM values for 500bp windows centered on the top 7,000 predicted target sites for the “sgRNA #1” in two *in vitro* CasKAS experiments. Off-target sites are color-coded by the number of mismatches relative to the sgRNA. (h) CasKAS requires a moderate sequencing depth of 10-20 10^6^ reads to accurately rank potential off-targets. (i-k) *In vitro* CasKAS maps Cas9 and dCas9 target sites. (i) Shown are CasKAS experiments with Cas9 and dCas9 and with the EMX1 sgRNA or with no sgRNA (negative control) (j) Assymmetric 5’ end distribution around target sites in dCas9 *in vivo* CasKAS. (k) In *in vivo* Cas9 CasKAS, a mixture distribution is observed between phased cleavage sites and broader ssDNA labeling.

To determine the feasibility of using KAS-seq to map regions of ssDNA generated by Cas9 binding, we carried out an initial *in vitro* experiment using mouse genomic DNA (gDNA), purified dCas9 and two sgRNAs targeting the *Hoxa* locus.

Strikingly, we observed strong peaks at the expected target sites for each sgRNA (Fig. 1c). Detailed examination of dCas9 CasKAS profiles around the predicted sgRNA target sites revealed strand coverage asymmetry patterns similar to those observed for ChIP-seq around transcription factor binding sites ^20^ (Fig. 1d), indicating that enrichment derives from the sgRNA target site itself and confirming the utility of N_3_-kethoxal for mapping dCas9 occupancy sites. We termed the assay “CasKAS”.

We then reasoned that CasKAS should also capture active Cas9 complexed with DNA, as the enzyme is thought to remain associated with DNA for some time after cleavage ^21^. We performed CasKAS experiments with the same sgRNAs and active Cas9 nuclease, and again observed enrichment at the expected on-target sites (Fig. 1e). Examination of Cas9 CasKAS read profiles around the on-target site showed that the 5’ ends of reads are precisely positioned around the expected cut site, with one cut position on the target strand (which binds the sgRNA and is cleaved by the HNH domain) and two to three such positions on the non-target strand (which is cleaved by the RuvC domain; Fig. 1f), consistent with the previously known patterns of Cas9 cleavage ^22,23^. CasKAS therefore provides target specificity profiles for both active and catalytically dead Cas9 enzymes.

*In vitro* CasKAS data was highly reproducible between replicates (Fig. 1g), and a modest sequencing depth of between 10 and 20 million mapped reads was sufficient to capture off-target specificity profiles (Fig. 1h), which is an order of magnitude lower than required for resequencing the whole genome.

We observed similar results with two mouse sgRNAs targeting the *Nanog* locus (Supplementary Fig. 1) and with two human sgRNA (“EMX1” and “VEGFA”; Supplementary Fig. 2 and 3). We found no enrichment using components of the RNP in isolation – sgRNAs, dCas9 or Cas9 (Supplementary Fig. 2).

Next we tested the application of CasKAS *in vivo*. Living cells contain substantial ssDNA due to active transcription and other processes ^19^, so *in vivo* CasKAS signal derives from a mixture of Cas9-associated ssDNA and endogenous processes. We carried out KAS-seq experiments using both dCas9 and Cas9 in HEK293 cells transfected with RNPs targeting *EMX1* or *VEGFA*, as well as negative, no-guide controls, which provided a map of background endogenous ssDNA profiles. At *EMX1*, which is not active in HEK293 cells, we observe strong peaks at the expected target site (Fig. 1i), as well as an asymmetric read profile around it for dCas9 (Fig. 1j), and a substantial degree of 5’ end clustering at the cut site, similar to what is observed *in vitro* for active Cas9 (Fig. 1g). The VEGFA gene is active in HEK293 cells, but the dCas9/Cas9 CasKAS signal is still readily identifiable as an addition to the endogenous ssDNA enrichment pattern (Supplementary Fig. 4). These results demonstrate the utility of CasKAS for profiling CRISPR specificity both *in vitro* and *in vivo*

We next examined the genome-wide specificity of sgR-NAs as measured by CasKAS. We focus on the mouse sgRNA #1 as it displayed a substantial number of off-targets yet that number was also sufficiently small for all of them to be examined directly. We first called peaks *de novo* (see Methods for details) without relying on off-target prediction algorithms, then manually curated the resulting peak set, excluding peaks not exhibiting the canonical asymmetric read distribution around a fixed point on the two strands (Fig. 2a). Remarkably, while we found 32 peaks at predicted off-target sites, we also found 198 (i.e. ∼6× as many) additional manually curated peaks; while these peaks exhibit generally lower CasKAS signal (Fig. 2b), they all display proper peak shape characteristics (see Supplementary Fig. 13 for details), suggesting that they are genuine sites of occupancy. Most of the predicted (in total ∼7,500) off-target sites for this sgRNA did not show substantial occupancy by dCas9 CasKAS (Fig. 2c-d).

**Figure 2:**
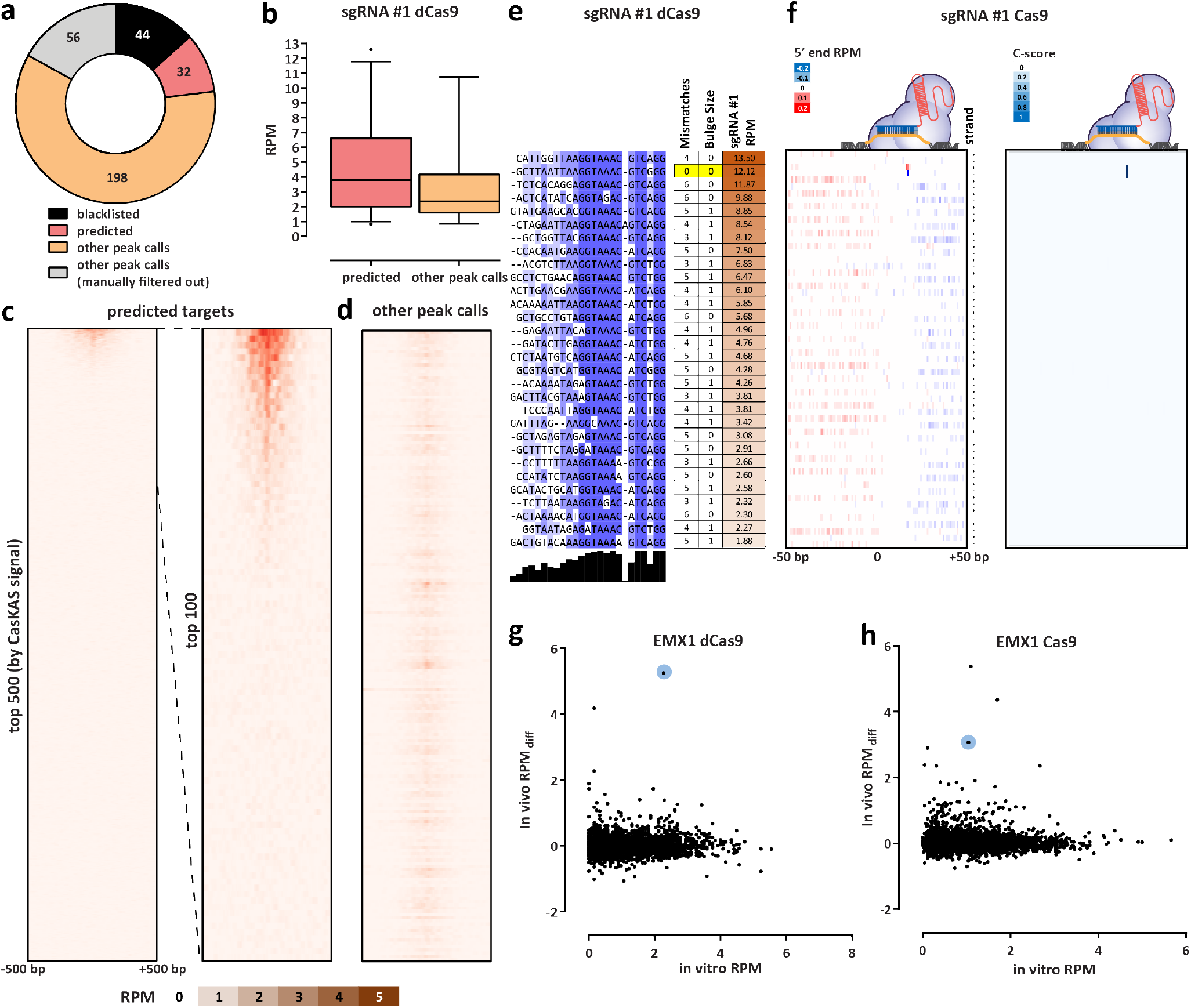
CasKAS profiles sgRNA specificity genome-wide. (a) Summary of de novo peak calls for sgRNA #1 (using MACS2) (b) CasKAS signal is stronger over predicted off-target sites, but legitimate interactions are also found elsewhere in the genome. (c) CasKAS profile over predicted (by Cas-OFFinder) off-target sites for sgRNA #1 with dCas9 (all such sites and focusing only on the top 100 ranked by dCas9 CasKAS signal). (d) CasKAS profile over peak calls outside predicted (by Cas-OFFinder) off-target sites for sgRNA #1 with dCas9. (e) Determinants of sequence specificity as measured by dCas9 CasKAS (for sgRNA #1). PAM-distal regions of the sgRNA are less constrained than its PAM-proximal parts. The on-target sgRNA is highlighted in yellow. (f) Active Cas9 signal read profiles can be used to distinguish off-targets associated with cutting from those where only binding occurs. Shown are the same off-target sites as in (e) and the plus- and minus-strand active Cas9 5’ end profiles around the sgRNA. In this case (sgRNA #1), only the on-target site shows a Cas9 CasKAS pattern indicating cleavage; at the other sites even active Cas9 likely only binds but does not cut. A simple cutting score metric (“*C*-score”) based on multiplying the 5’ end forward- and reverse-strand profiles can be used to quantify cutting vs. binding. (g and h) Comparison between *in vitro* and *in vivo* CasKAS signal over predicted off-target sites for the EMX1 sgRNA. *In vivo* CasKAS is quantified as the difference in read per million (±500 bp of the sgRNA site) between the sgRNA KAS-seq and the no-guide control KAS-seq (“RPM_*diff*_). The on-target site is shown in blue.

Sequence comparison of the occupied predicted off-target sites allowed us to evaluate determinants of Cas9 specificity (Fig. 2e). Consistent with previous reports ^24,25^, the PAM-distal region was much less sequence-constrained than the PAM-proximal seed region. We observed a similar pattern with the other sgRNAs we profiled, in both mouse and human (Supplementary Fig. 5-8 and Supplementary Fig. 9-12).

When analyzing peaks not associated with predicted off-target sites (Supplementary Fig. 14) we observed other telling patterns – at numerous sites with strong dCas9 CasKAS signal, we observe a large number of mismatches to the sgRNA sequence as well as “bulge” regions wherein indels are observed in the target sequence. These mismatches and bulges were in general much larger than what is considered permissible by off-target prediction algorithms; we speculate that the lack of consideration of potential target sequences with large numbers of mismatches or substantial insertions could explain the much larger number of such sites compared to the set of occupied predicted off-targets.

We next devised a simple metric for evaluating the degree of read clustering at cut sites (a “*C*-score”; see Methods for details) to estimate the degree of cutting by Cas9. The on-target site exhibits the second highest dCas9 CasKAS signal genome-wide. However, strikingly, even though all CasKAS-identified off-target sites showed Cas9 binding, only the on-target site displayed strong cutting activity (Fig. 2f). The behavior of other sgRNAs varies (Supplementary Fig. 5-8 and 15), with some showing multiple clearly identifiable cut sites. Overall, these results are consistent with previous reports that Cas9 requires more successful RNA:DNA basepairing for cleavage activity than is necessary for binding ^26,27^. Thus, interpreting the read distributions of Cas9 CasKAS at target sites enables simultaneous detection of binding specificity and the promiscuity of catalytic activity.

Finally, we compared *in vitro* and *in vivo* CasKAS profiles (Fig. 2g-h). We find many fewer strongly enriched sites in *in vivo* datasets than *in vitro*, with the on-target site being either the top (for dCas9) or among the top (for Cas9) sites in vivo. A potential explanation for this difference is the previously reported impediment of Cas9/dCas9 binding to DNA by the presence of nucleosomes ^28^.

In conclusion, we have presented CasKAS, a simple and robust method for mapping the specificity of active and catalytically dead versions of CRISPR enzymes. CasKAS has numerous advantages over existing tools while also opening up new possibilities for studying CRISPR biology. CasKAS requires no specialized molecular biology protocols, takes just a few hours *in vitro* (and a similar amount of time after harvesting cells *in vivo*), and, due to the strong, active enrichment of target sequences, is inexpensive. In contrast to previously developed methods, It measures strand invasion by CRISPR, which is biochemically more specific and relevant to CRISPR function than DNA association. We compared *de novo* called CasKAS peaks to those generated by other means, and while we found large sets of peaks unique to each method, those found only by CasKAS contained much higher fractions of predicted off-target sites than those unique to other methods (Supplementary Fig. 16).

CasKAS does not rely on measuring DNA cleavage or modification and can thus be used to profile the specificity of all types of DNA-targeting CRISPR proteins. CasKAS also does not rely on cellular repair processes, cell division, or delivery of additional exogenous DNA (as in GUIDE-seq) to generate a detectable signal. These advantages, coupled with low cell input requirements, may increase the utility of the method in rare primary cell types, tissues from animal models, or even for direct assessment of specificity in edited patient cells (e.g. *ex vivo* edited immune cells). A current limitation of CasKAS is the requirement that a G nucleotide is present within the sgRNA sequence, since kethoxal requires an exposed G to react with. However, only a small fraction (≤5%) of sgRNAs in the human genome lack any Gs for *S. pyogenes* PAM sequences (Supplementary Fig. 17). Another minor limitation of the current *in vitro* protocol is that labeling is carried out on high molecular weight (HMW) DNA and samples must be sheared serially. We have explored using pre-sheared and end-repaired DNA (to minimize kethoxal labeling of Gs on sticky ends generated by sonication), with comparable results to using HMW DNA (Supplementary Fig. 18); we anticipate that further optimization or using other approaches, such as enzymatic fragmentation, should allow the parallel high-throughput plate-based profiling of the specificity of very large numbers of sgRNAs.

In addition to being highly valuable for off-target profiling *in vitro* and in previously difficult to assay settings such as primary cells, we expect CasKAS to provide fruitful insights into the mechanisms and dynamics of *in vivo* CRISPR action (taking advantage of finely controllable CRISPR systems such as vfCRISPR ^29^), and the influence of transcriptional, regulatory, and epigenetic and other functional genomic contexts on CRISPR activity.

## Methods

### Guide RNA sequences

Guide RNAs were obtained from IDT (“sgRNA #1” and “sgRNA #2”) or from Synthego (all others).

The following sgRNA sequences were used in this study:

1. “sgRNA #1”: GCTTAATTAAGGTAAACGTC
2. “sgRNA #2”: CCAACCTGGCGGCTCGTTGG
3. “EMX1 Tsai”: GAGTCCGAGCAGAAGAAGAA
4. “VEGFA-site1”: GGGTGGGGGGAGTTTGCTCC
5. “Nanog-sg2”: GATCTCTAGTGGGAAGTTTC
6. “Nanog-sg3”: GTCTGTAGAAAGAATGGAAG

Guide RNAs were dissolved to a concentration of 100 *µ*M using nuclease-free 1× TE buffer and stored at –20 ^*°*^C.

### *In vitro* CasKAS

*In vitro* CasKAS experiments were executed as follows.

First, 1 *µ*L of each synthetic sgRNA were incubated at room temperature with 1 *µ*L of recombinant purified dCas9 (MCLab dCAS9B-200) for 20 minutes. The RNP was then incubated with 1 *µ*g of gDNA at 37 ^*°*^C for 10 minutes.

The KAS reaction was then carried out by adding 1 *µ*L of 500 mM N_3_-kethoxal (ApeXBio A8793). DNA was immediately purified using the MinElute PCR Purification Kit (Qiagen 28006), and eluted in 87.5 or 175 *µ*L 25mM K_3_BO_3_.

### *In vivo* CasKAS

For *in vivo* CasKAS experiments, HEK293T cells were seeded at 400,000 cells/well into a 6-well plate the day before RNP transfection. Media was exchanged 2 hours before transfection. For each well, 6,250 ng of Cas9 (MCLAB CAS9-200) or dCas9 (MCLAB dCAS9B-200) and 1,200 ng sgRNA was complexed with CRISPRMAX (Thermo Fisher CMAX00008) reagent in Opti-MEM (Thermo Fisher 51985091) following manufacturer’s protocol. After incubation at room temperature for 15 minutes, the RNP solution was directly added to each well and gently mixed. The cells were incubated with the RNP complex for 14 hours at 37 ^*°*^C. To harvest and perform kethoxal labeling, media was removed and room temperature 1×PBS was used to wash the cells. Cells were then dissociated with trypsin, trypsin was quenched with media, cells were pelleted at room temperature, and then resuspended in 100 *µ*L of media supplemented with 5 M N_3_-kethoxal (final concentration). Cells were incubated for 10 minutes at 37 ^*°*^C with shaking at 500 rpm in a Thermomixer. Cells were then pelleted by centrifuging at 500 *g* for 5 minutes at 4 ^*°*^C. Genomic DNA was then extracted using the Monarch gDNA Purification Kit (NEB T3010S) following the standard protocol but with elution using 175 *µ*L 25 mM K_3_BO_3_ at pH 7.0.

### Click reaction, biotin pull down and library generation

The click reaction was carried out by combining 175 *µ*L purified DNA, 5 *µ*L 20 mM DBCO-PEG4-biotin (DMSO solution, Sigma 760749), and 20 *µ*L 10×PBS in a final volume of 200 *µ*L or 87.5 *µ*L purified and sheared DNA, 2.5 *µ*L 20 mM DBCO-PEG4-biotin (DMSO solution, Sigma 760749), and 10 *µ*L 10×PBS in a final volume of 100 *µ*L. The reaction was incubated at 37 ^*°*^C for 90 minutes.

DNA was purified using AMPure XP beads (50 *µ*L for a 100 *µ*L reaction or 100 *µ*L for a 200 *µ*L reaction), beads were washed on a magnetic stand twice with 80% EtOH, and eluted in 130 *µ*L 25mM K_3_BO_3_.

Purified DNA was then sheared on a Covaris E220 instrument down to ∼150-400 bp size.

For streptavidin pulldown of biotin-labeled DNA, 10 *µ*L of 10 mg/mL Dynabeads MyOne Streptavidin T1 beads (Life Technologies, 65602) were separated on a magnetic stand, then washed with 300 *µ*L of 1×TWB (Tween Washing Buffer; 5 mM Tris-HCl pH 7.5; 0.5 mM EDTA; 1 M NaCl; 0.05% Tween 20). The beads were resuspended in 300 *µ*L of 2×Binding Buffer (10 mM Tris-HCl pH 7.5, 1 mM EDTA; 2 M NaCl), the sonicated DNA was added (diluted to a final volume of 300 *µ*L if necessary), and the beads were incubated for≥15 minutes at room temperature on a rotator. After separation on a magnetic stand, the beads were washed with 300 *µ*L of 1×TWB, and heated at 55 ^*°*^C in a Thermomixer with shaking for 2 minutes. After removal of the supernatant on a magnetic stand, the TWB wash and 55 ^*°*^C incubation were repeated.

Final libraries were prepared on beads using the NEB-Next Ultra II DNA Library Prep Kit (NEB, #E7645) as follows. End repair was carried out by resuspending beads in 50 *µ*L 1×EB buffer, and adding 3 *µ*L NEB Ultra End Repair Enzyme and 7 *µ*L NEB Ultra End Repair Enzyme, followed by incubation at 20 ^*°*^C for 30 minutes (in a Thermomixer, with shaking at 1,000 rpm) and then at 65 ^*°*^C for 30 minutes.

Adapters were ligated to DNA fragments by adding 30 *µ*L Blunt Ligation mix, 1 *µ*L Ligation Enhancer and 2.5 *µ*L NEB Adapter, incubating at 20 ^*°*^C for 20 minutes, adding 3 *µ*L USER enzyme, and incubating at 37 ^*°*^C for 15 minutes (in a Thermomixer, with shaking at 1,000 rpm).

Beads were then separated on a magnetic stand, and washed with 300 *µ*L TWB for 2 minutes at 55 ^*°*^C, 1000 rpm in a Thermomixer. After separation on a magnetic stand, beads were washed in 100 *µ*L 0.1×TE buffer, then resuspended in 15 *µ*L 0.1×TE buffer, and heated at 98 ^*°*^C for 10 minutes.

For PCR, 5 *µ*L of each of the i5 and i7 NEB Next sequencing adapters were added together with 25 *µ*L 2×NEB Ultra PCR Mater Mix. PCR was carried out with a 98 ^*°*^C incubation for 30 seconds and 12 cycles of 98 ^*°*^C for 10 seconds, 65 ^*°*^C for 30 seconds, and 72 ^*°*^C for 1 minute, followed by incubation at 72 ^*°*^C for 5 minutes.

Beads were separated on a magnetic stand, and the supernatant was cleaned up using 1.8× AMPure XP beads.

Libraries were sequenced in a paired-end format on an Illumina NextSeq instrument using NextSeq 500/550 high output kits (2×36 cycles).

### Data processing

Demultipexed fastq files were mapped to the `hg38` assembly of the human genome or the `mm10` version of the mouse genome as 2×36mers using Bowtie ^30^ with the following settings: `-v 2 -k 2 -m 1 --best --strata -X 1000`. Duplicate reads were removed using `picard-tools` (version 1.99).

Browser tracks generation, fragment length estimation, TSS enrichment calculations, and other analyses were carried out using custom-written Python scripts (https://github.com/georgimarinov/GeorgiScripts). The refSeq set of annotations were used for evaluation of enrichment around TSSs.

### Peak calling

Peak calling on *in vitro* binding datasets was carried out using version 2.1.0 of MACS2^31^ with default settings.

Peaks were then compared against the ENCODE set of “blacklisted” regions ^32^ to filter out likely artifacts.

### Sequence analysis

Guide RNA off-target predictions were obtained from Cas-OFFinder ^33^

Multiple sequence alignments of sgRNA sequences and their off-targets were generated using MUSCLE ^34^ and visualized using JalView ^35^.

### Quantification

#### Cutting score calculation

The Cas9 cutting *C*-score was calculated as follows.

First, basepair-level Read-Per-Million (RPM) profiles for mapped read 5’ ends were generated separately for the forward and reverse strands. Then the *C*-score was calculated by multiply the forward and reverse strand profiles (summed over a running window of 3 bp):

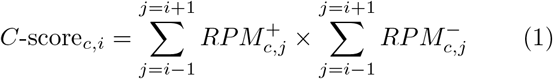

Where *c, i* indicate the coordinates by chromosome and position.

## Data availability

Sequencing reads for the datasets described in this study are available from GEO accession GSE171962.

## Author contributions

G.K.M. conceptualized the study, performed initial *in vitro* CasKAS experiments, analyzed data, and wrote the manuscript with input from all authors. S.H.K. developed the *in vivo* CasKAS protocol and performed *in vivo* CasKAS experiments. S.T.B. carried out *in vitro* CasKAS optimization. A.E.T. and J.T. supplied sgRNAs and designed off-target profiling experiments. A.E.T. carried out off-target analysis for mouse sgRNAs. T.W. provided key reagents. W.J.G., A.K., C.H. M.C.B. and L.B. supervised the study.

## Acknowledgments

This work was supported by NIH grants (P50HG007735, RO1 HG008140, U19AI057266 and UM1HG009442 to W.J.G., 1UM1HG009436 to W.J.G. and A.K., 1DP2OD022870-01 and 1U01HG009431 to A.K., and HG006827 to C.H.), the Rita Allen Foundation (to W.J.G.), the Baxter Foundation Faculty Scholar Grant, and the Human Frontiers Science Program grant RGY006S (to W.J.G). W.J.G is a Chan Zuckerberg Biohub investigator and acknowledges grants 2017-174468 and 2018-182817 from the Chan Zuckerberg Initiative. S.K. is supported by MSTP training grant T32GM007365 and the Paul and Daisy Soros Fellowship. J.T. is supported by the NIDDK F99/K00 fellowship of the National Institutes of Health (F99DK126120). M.C.B. is supported by a grant from Stanford ChEM-H and an NIH Directors New Innovator Award (1DP2HD08406901). Fellowship support also provided by the Stanford School of Medicine Dean’s Fellowship (G.K.M.), the Siebel Scholars, the Enhancing Diversity in Graduate Education Program and the Weiland Family Fellowship (A.E.T.). C.H. is a Howard Hughes Medical Institute Investigator.

The authors would like to thank Zohar Shipony and members of the Greenleaf, Kundaje, and Bassik labs for helpful discussion and suggestions regarding this work.

## Supplementary Materials

### Supplementary Figures

**Supplementary Figure 1:**
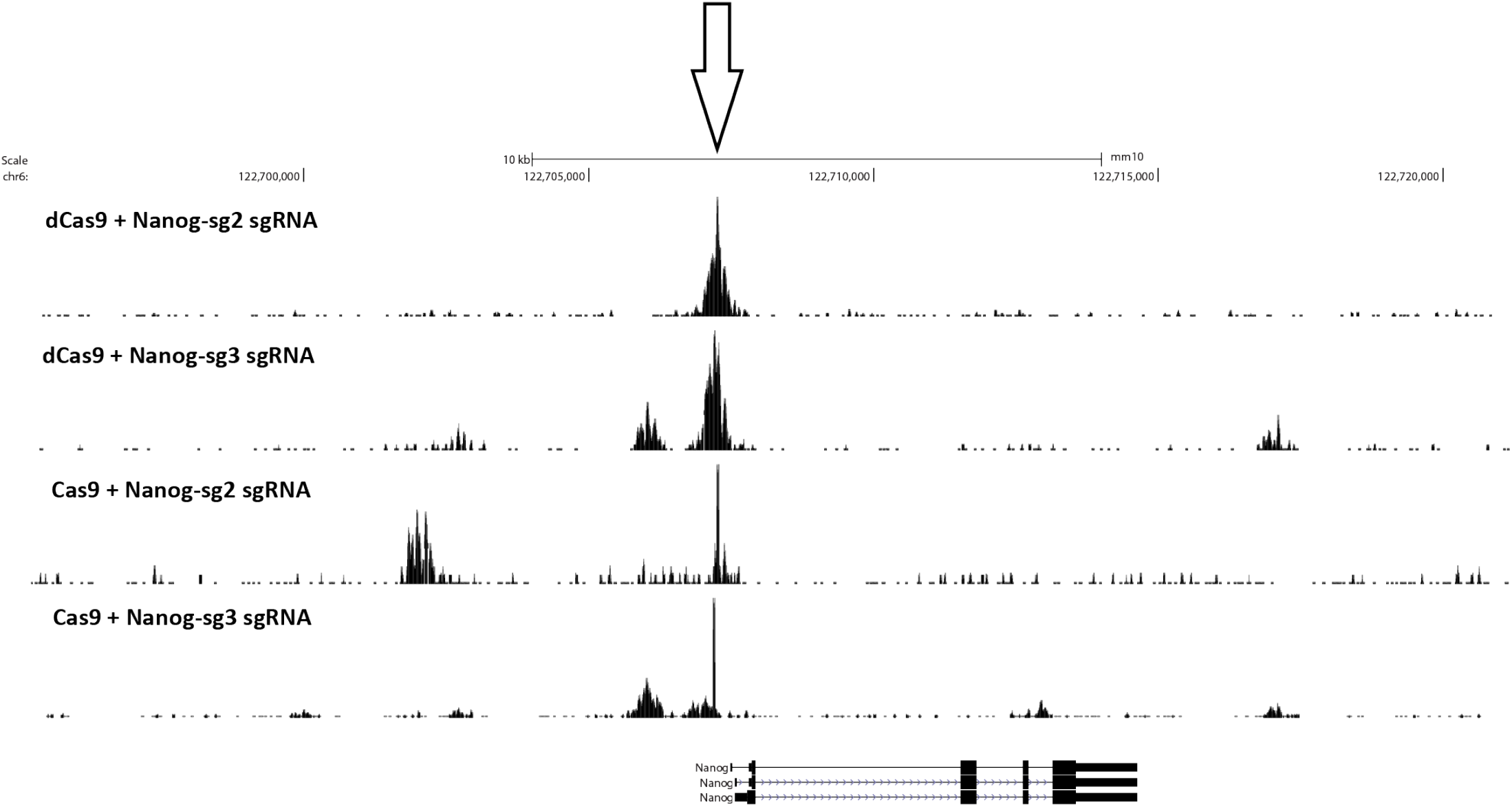
*In vitro* dCas9 and Cas9 CasKAS profiles around the mouse *Nanog* locus using the “Nanog-sg2” and “Nanog-sg3” sgRNAs.

**Supplementary Figure 2:**
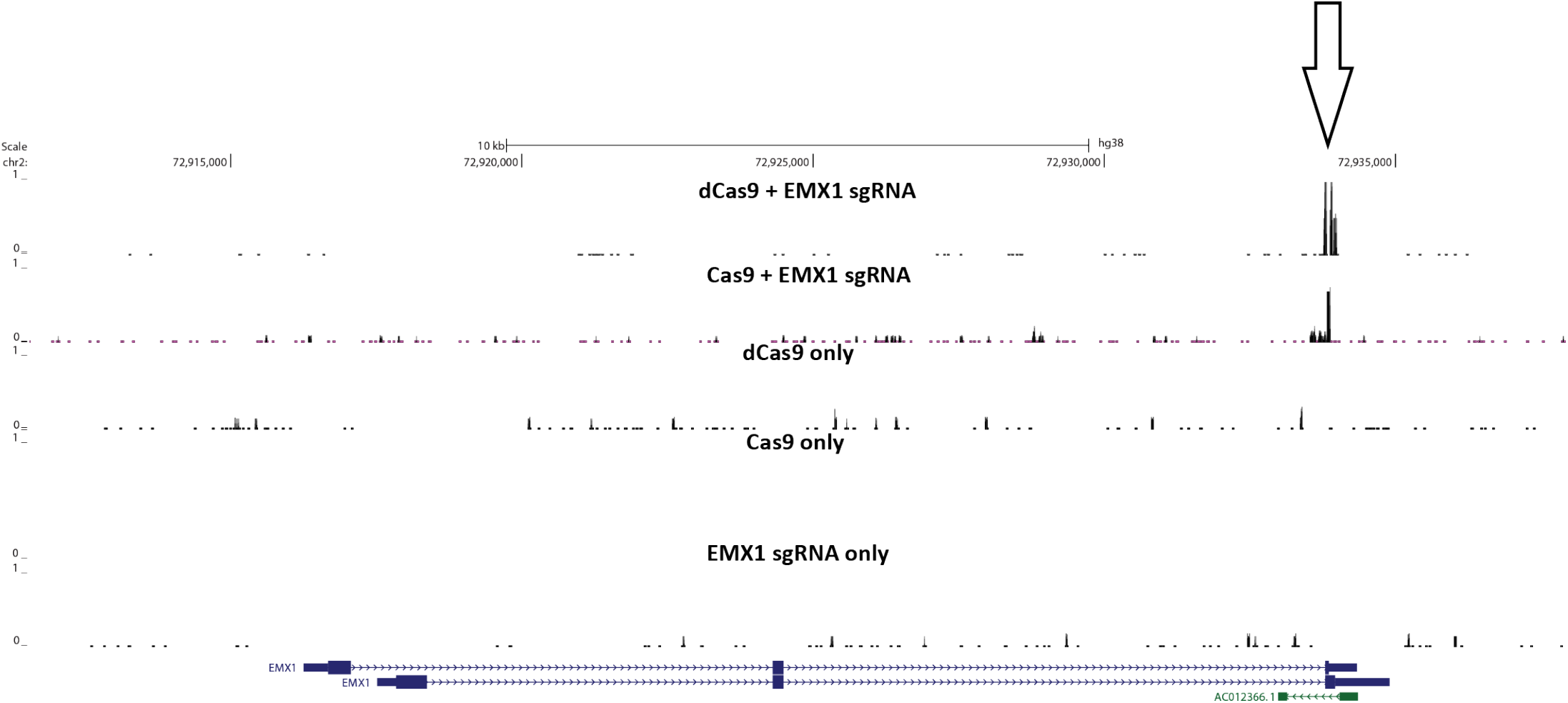
CasKAS signal *in vitro* is specific to the activity of the dCas9/Cas9 protein combined with its sgRNA. CasKAS was carried out with the EMX1 sgRNA and with the following combinations of protein and sgRNA: dCas9 + sgRNA, Cas9 + sgRNA, dCas9 alone, Cas9 alone, or sgRNA alone.

**Supplementary Figure 3:**
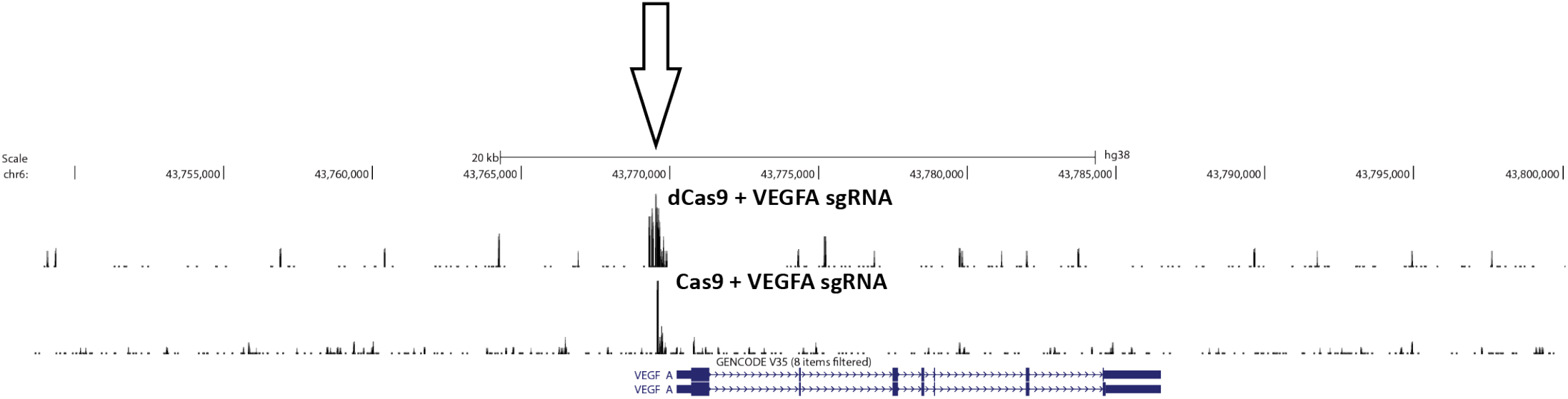
CasKAS signal *in vitro* around the *VEGFA* gene with the VEGFA sgRNA.

**Supplementary Figure 4:**
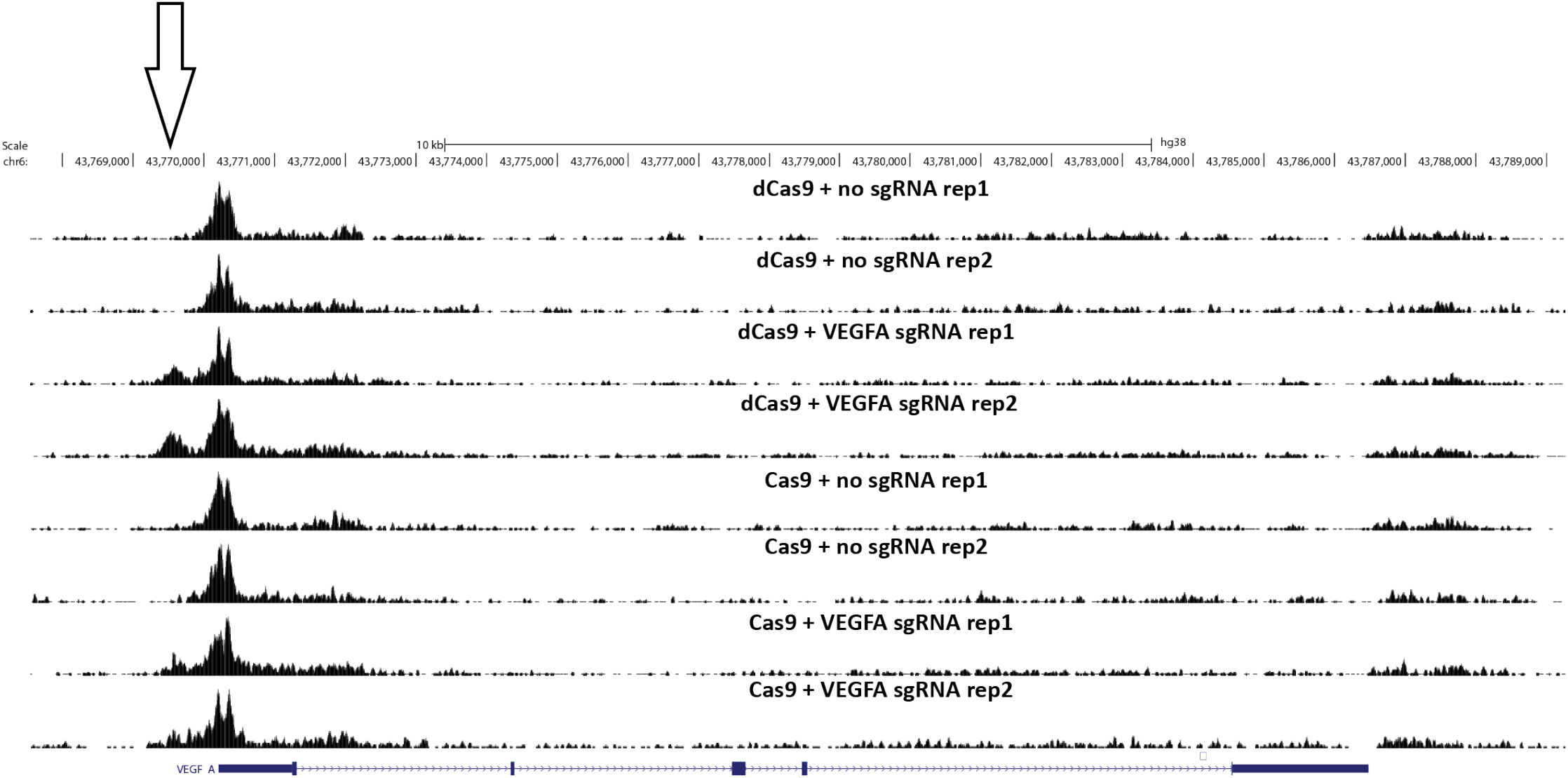
CasKAS signal *in vivo* around the *VEGFA* gene with the VEGFA sgRNA.

**Supplementary Figure 5:**
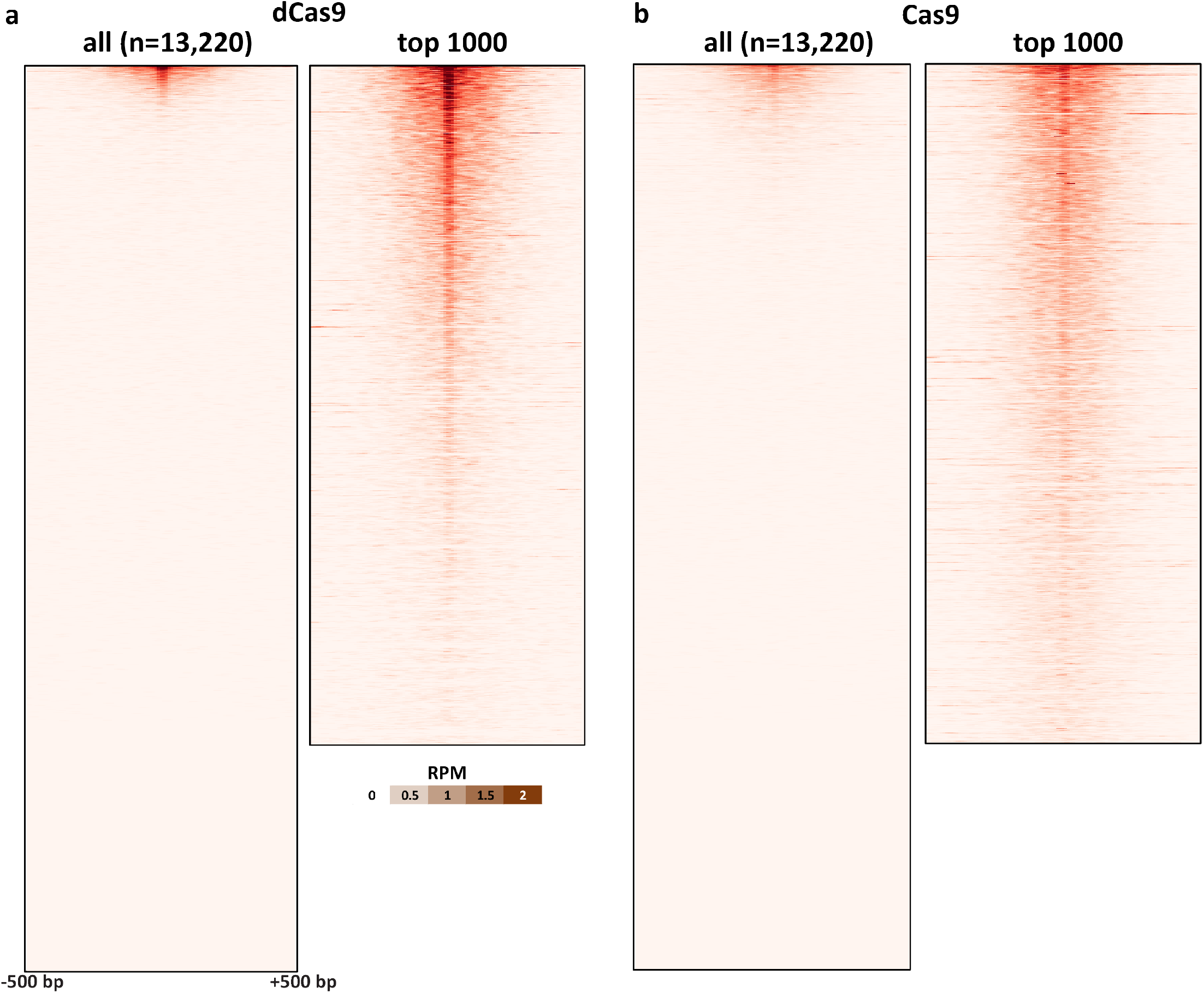
*In vitro* dCas9 and Cas9 CasKAS profiles for the “Nanog-sg2” sgRNA. CasKAS profiles are shown for all off-target sites predicted by Cas-OFFinder as well as for the top 1000 sites (ranked by CasKAS RPM values over the ±500bp region around the sgRNA target site).

**Supplementary Figure 6:**
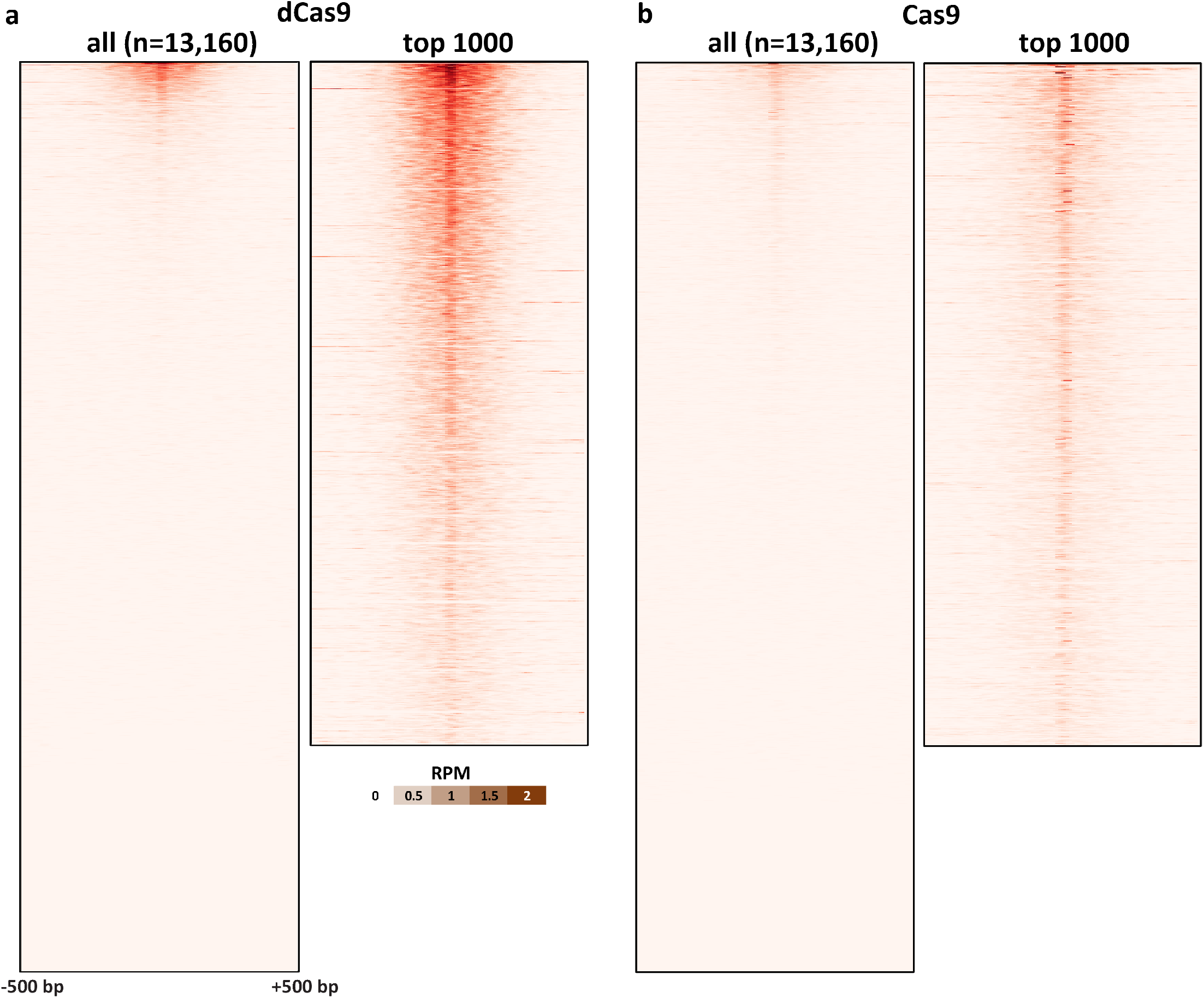
*In vitro* dCas9 and Cas9 CasKAS profiles for the “Nanog-sg3” sgRNA. CasKAS profiles are shown for all off-target sites predicted by Cas-OFFinder as well as for the top 1000 sites (ranked by CasKAS RPM values over the ±500bp region around the sgRNA target site).

**Supplementary Figure 7:**
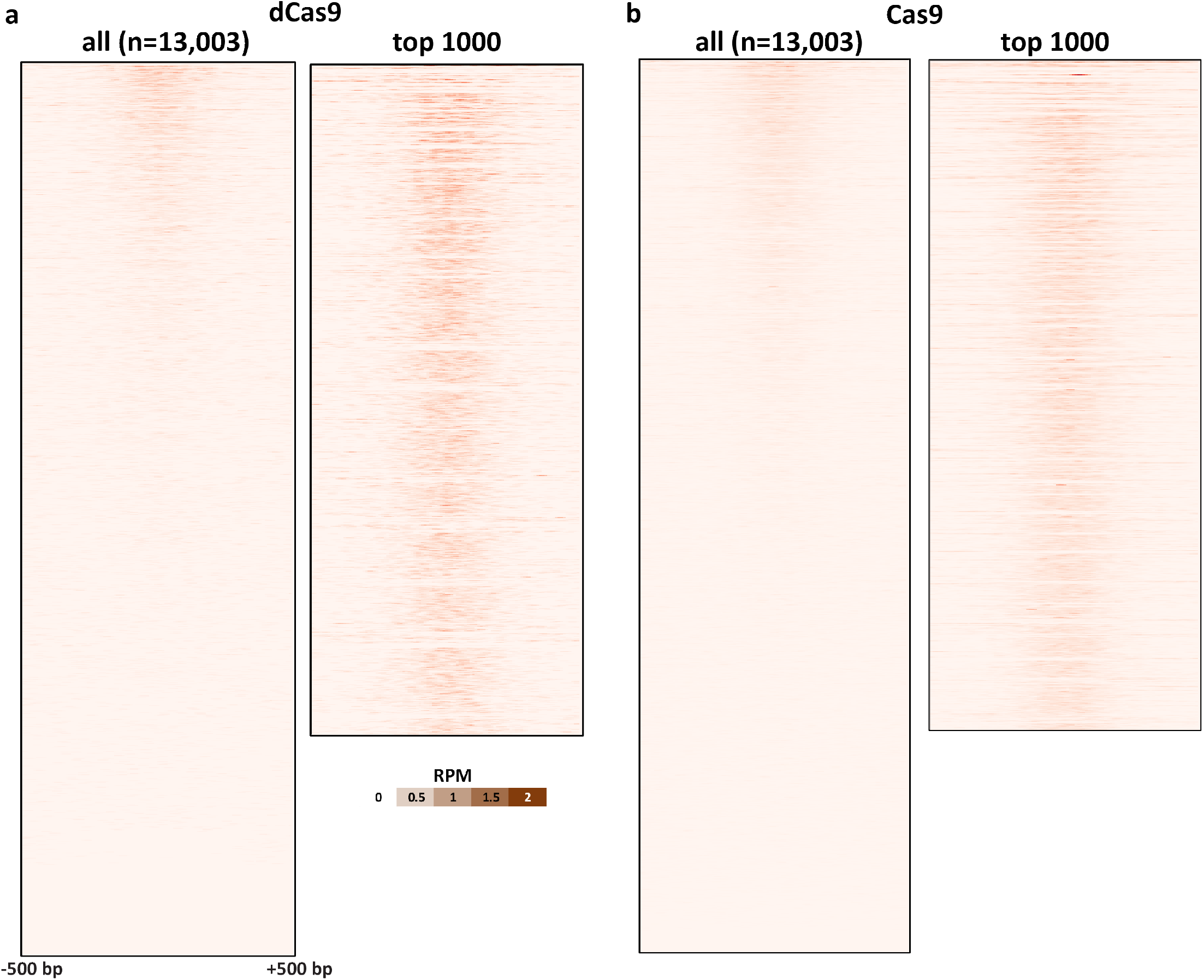
*In vitro* dCas9 and Cas9 CasKAS profiles for the “EMX1 Tsai” sgRNA. CasKAS profiles are shown for all off-target sites predicted by Cas-OFFinder as well as for the top 1000 sites (ranked by CasKAS RPM values over the ±500bp region around the sgRNA target site).

**Supplementary Figure 8:**
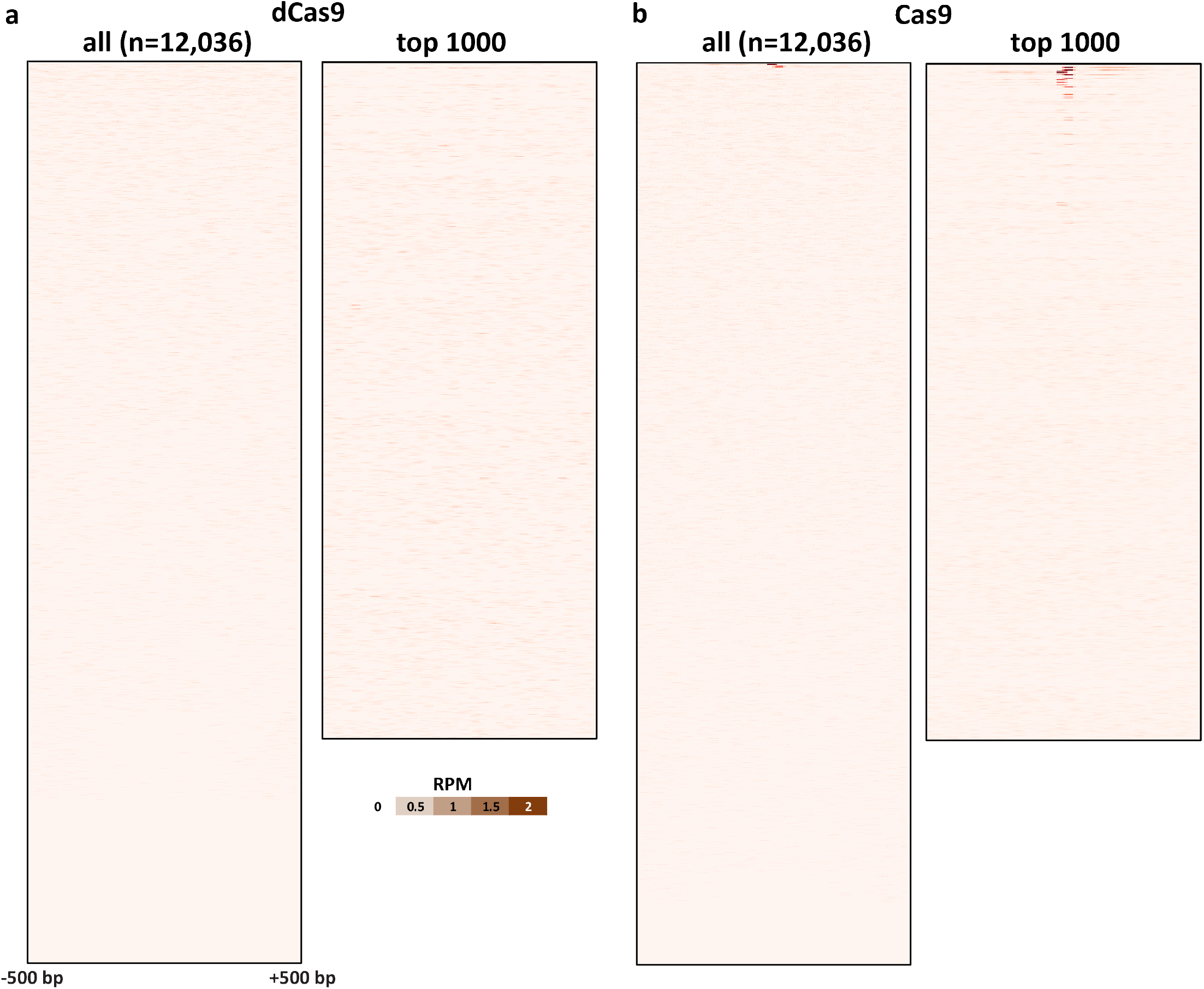
*In vitro* dCas9 and Cas9 CasKAS profiles for the “VEGFA-site1” sgRNA. CasKAS profiles are shown for all off-target sites predicted by Cas-OFFinder as well as for the top 1000 sites (ranked by CasKAS RPM values over the ±500bp region around the sgRNA target site).

**Supplementary Figure 9:**
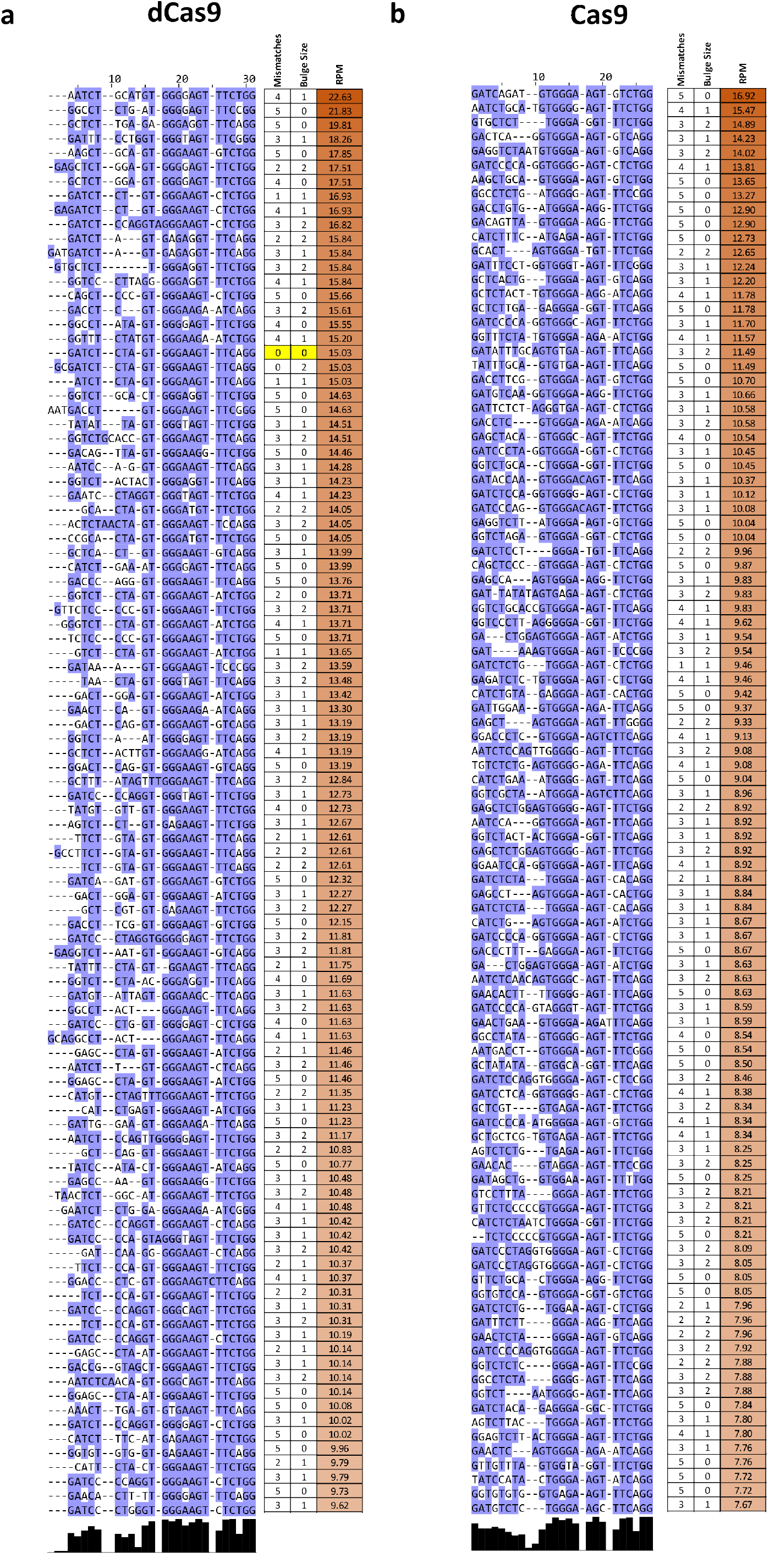
Multiple sequence alignment of off-target sites identified by *in vitro* dCas9 and Cas9 CasKAS for the “Nanog-sg2” sgRNA. Shown are the top 100 off-target sites as predicted by Cas-OFFinder and ranked by CasKAS signal. The on-target site (if within the top 100) is highlighted in yellow.

**Supplementary Figure 10:**
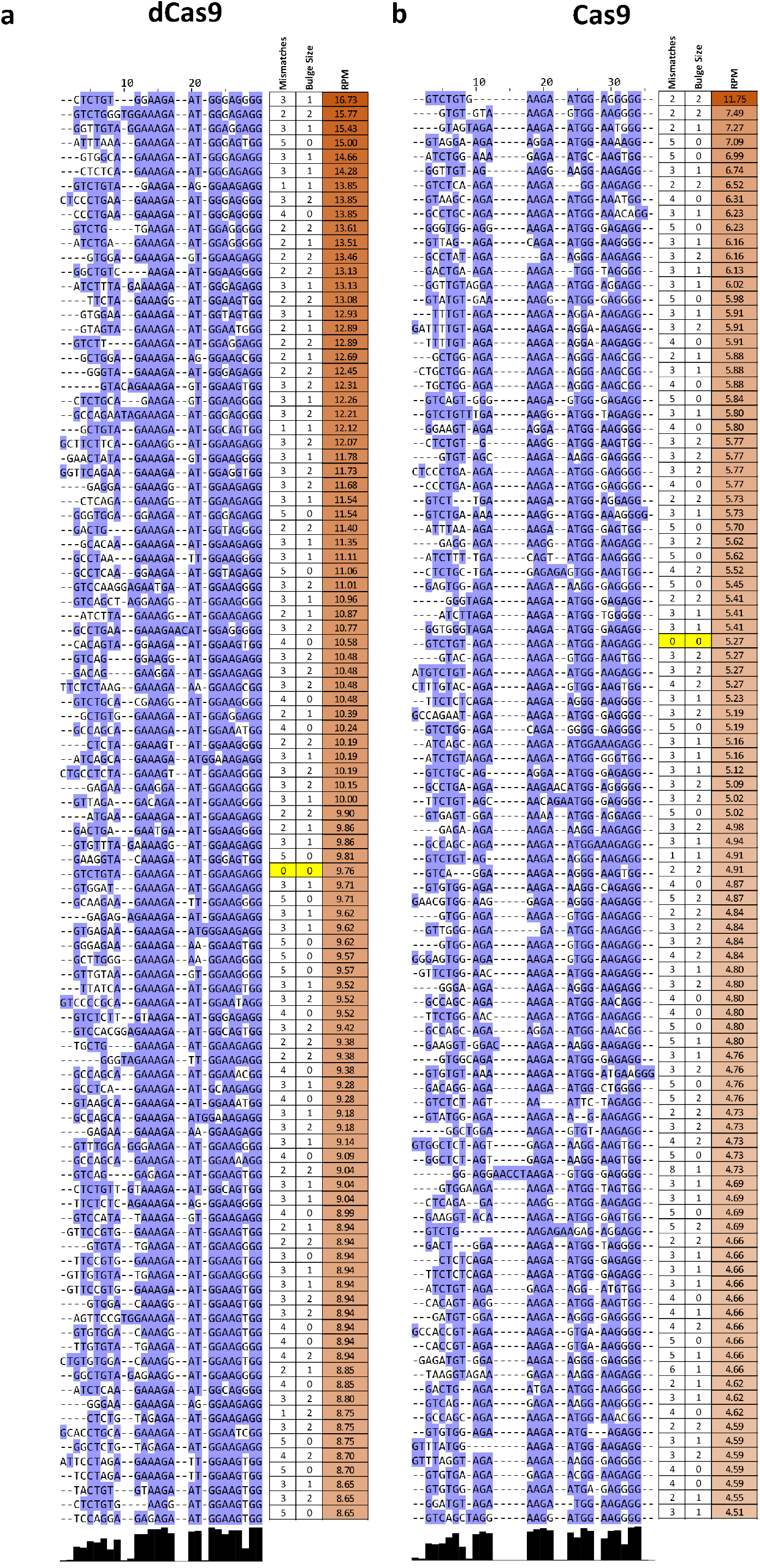
Multiple sequence alignment of off-target sites identified by *in vitro* dCas9 and Cas9 CasKAS for the “Nanog-sg3” sgRNA. Shown are the top 100 off-target sites as predicted by Cas-OFFinder and ranked by CasKAS signal. The on-target site (if within the top 100) is highlighted in yellow.

**Supplementary Figure 11:**
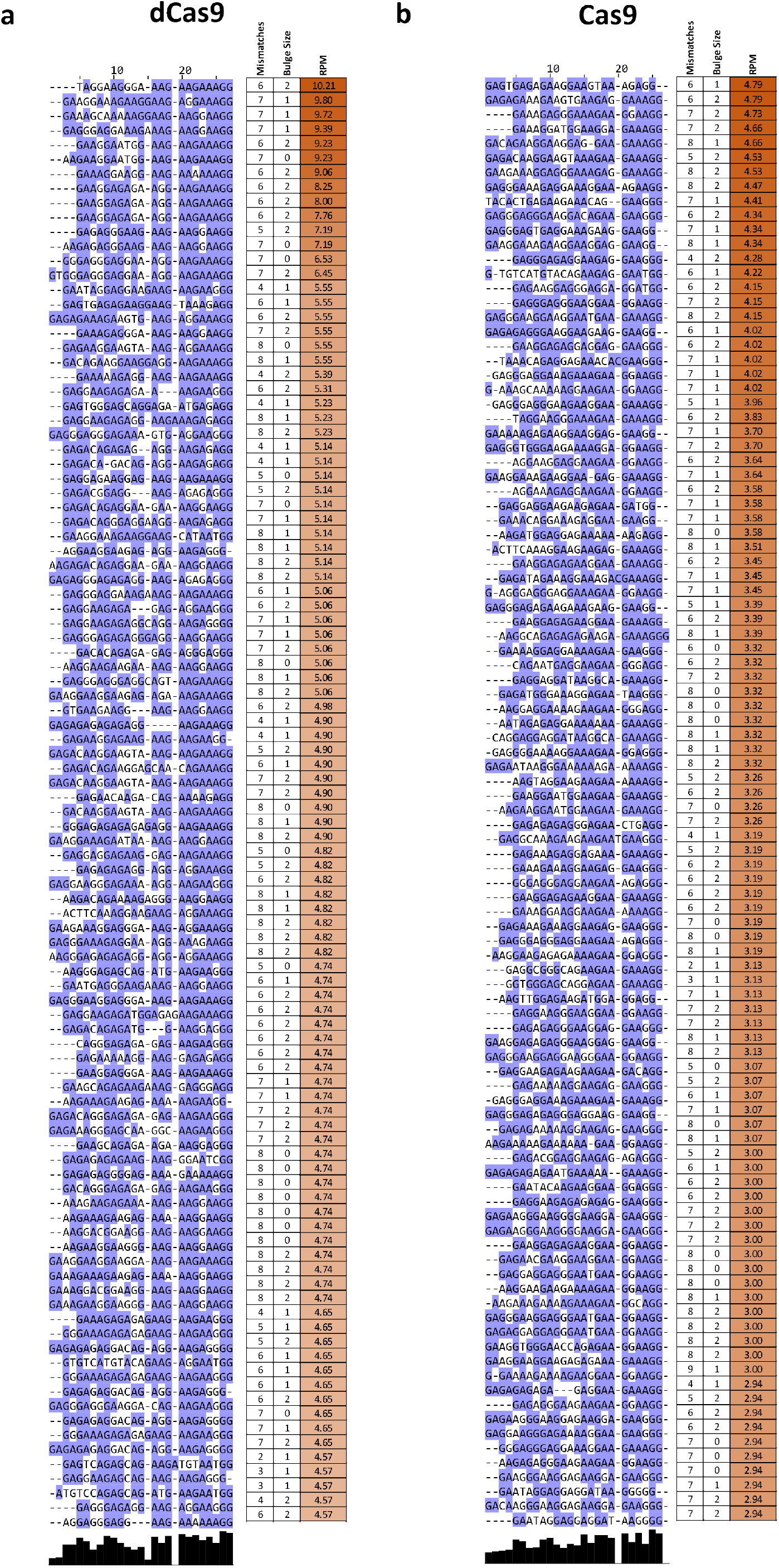
Multiple sequence alignment of off-target sites identified by *in vitro* dCas9 and Cas9 CasKAS for the “EMX1 Tsai” sgRNA. Shown are the top 100 off-target sites as predicted by Cas-OFFinder and ranked by CasKAS signal. The on-target site (if within the top 100) is highlighted in yellow.

**Supplementary Figure 12:**
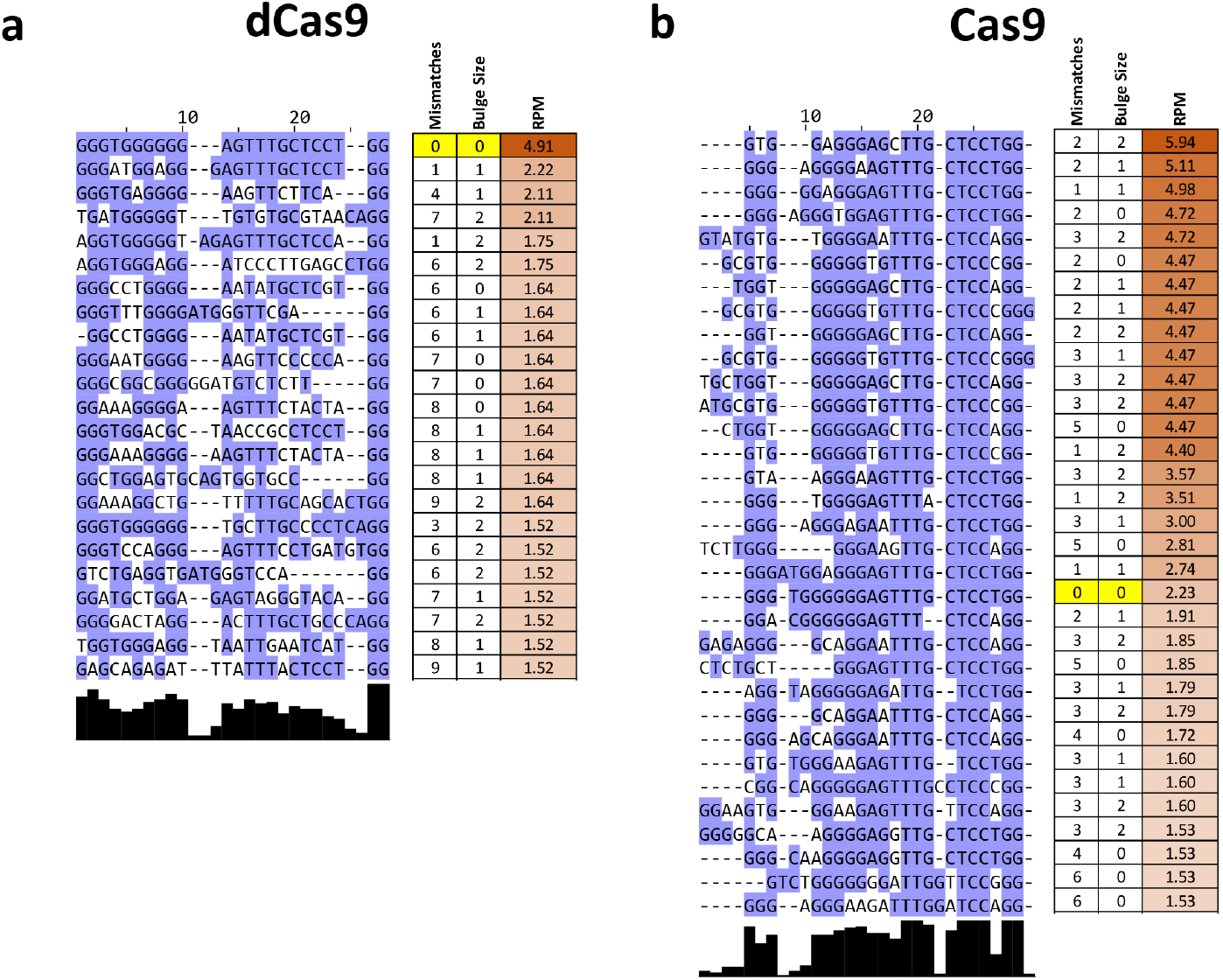
Multiple sequence alignment of off-target sites identified by *in vitro* dCas9 and Cas9 CasKAS for the “VEGFA-site1” sgRNA. Shown are the all target sites with RPM ≥1.5 as predicted by Cas-OFFinder and ranked by CasKAS signal. The on-target site (if within the top 100) is highlighted in yellow.

**Supplementary Figure 13:**
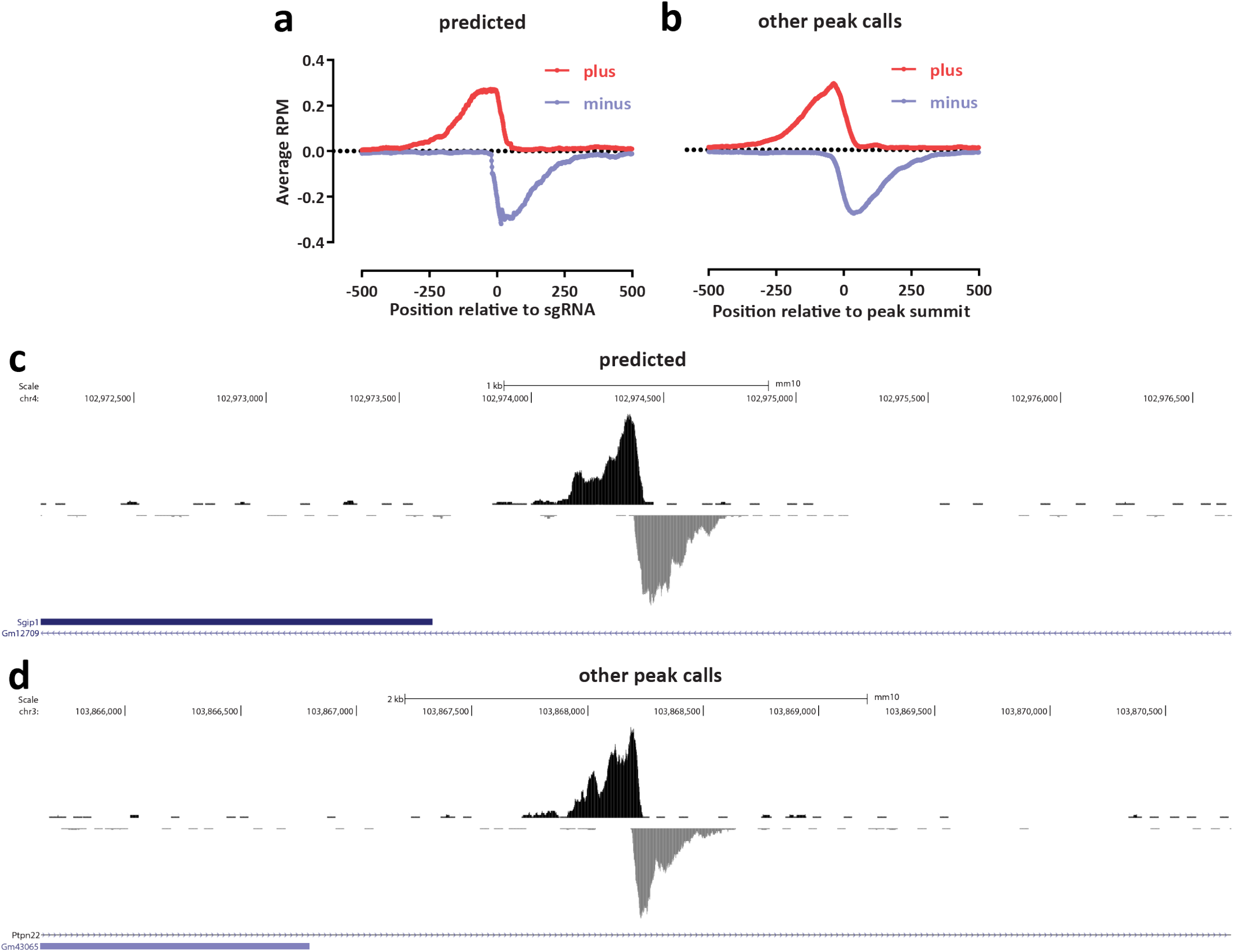
CasKAS identifies proper off-target sites that are missed by sgRNA prediction algorithms. Shown is *in vitro* dCas9 CasKAS for the “sgRNA #1” sgRNA. Peaks were called *de novo* using MACS2, then intersected with Cas-OFFinder off-target prediction, and the outersect was manually filtered to exclude obvious artifacts based on peak shape (e.g. arising from repetitive elements in the genome). (a) Aggregate forward- and reverse-strand profiles around off-target sites predicted by Cas-OFFinder (centered on the sgRNA); (b) Aggregate forward- and reverse-strand profiles around sites not predicted by Cas-OFFinder (centered on the MACS2 peak summit); (c) Example UCSC Genome Browser snapshot of a CasKAS read profile around an off-target site predicted by Cas-OFFinder; (c) Example UCSC Genome Browser snapshot of a CasKAS read profile around an off-target site not predicted by Cas-OFFinder. Both predicted and identified through peak calling sites exhibit the expected asymmetric read distribution around a fixed occupancy point (the sgRNA-dCas9 RNP complexed with DNA).

**Supplementary Figure 14:**
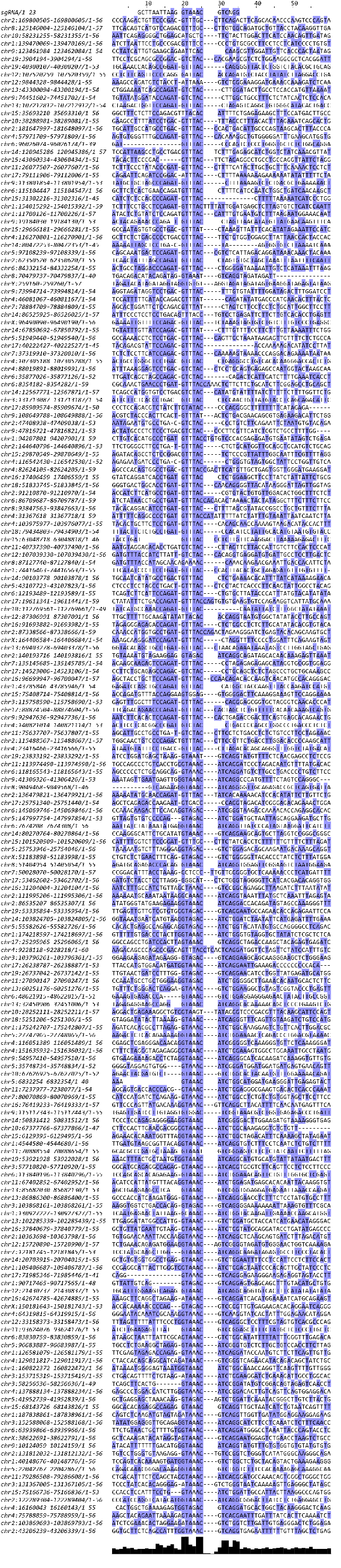
Multiple sequence alignment of off-target sites identified by *in vitro* dCas9 and Cas9 CasKAS for the “sgRNA #1” sgRNA outside the list of predicted off-targets by Cass-OFFinder. MACS2 peak calls were manually filtered to exclude artifactual peaks, then the sequence of the ±50-bp region around the peak summit was used as input to the multiple sequence alignment, together with the sgRNA itself.

**Supplementary Figure 15:**
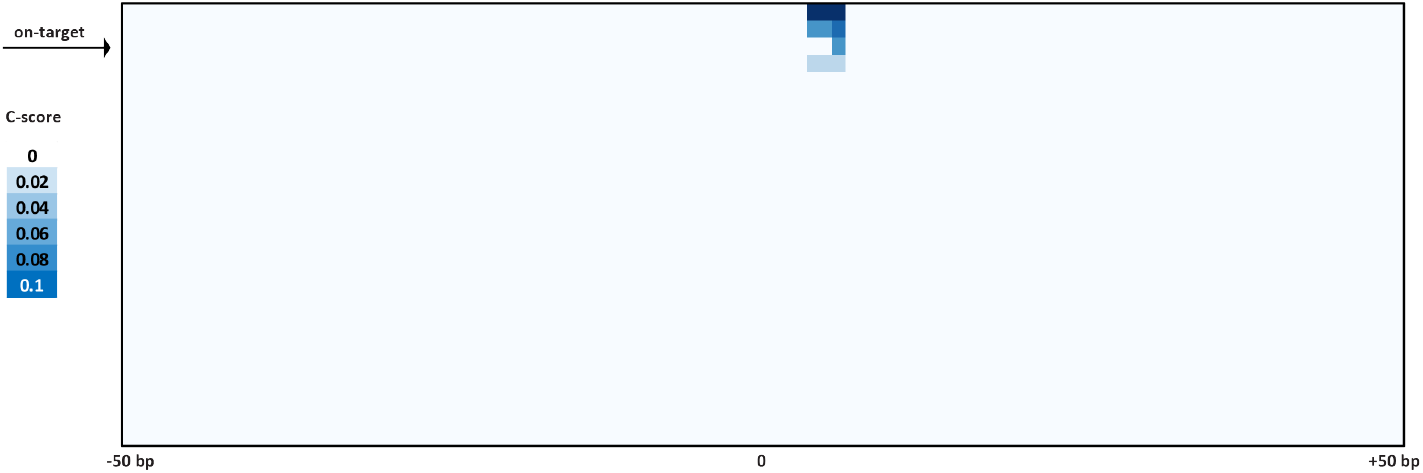
Cutting profiles around on- and off-target sites for the VEGFA sgRNA. Four sites where cleavage is observed are identified within the list of predicted off-targets.

**Supplementary Figure 16:**
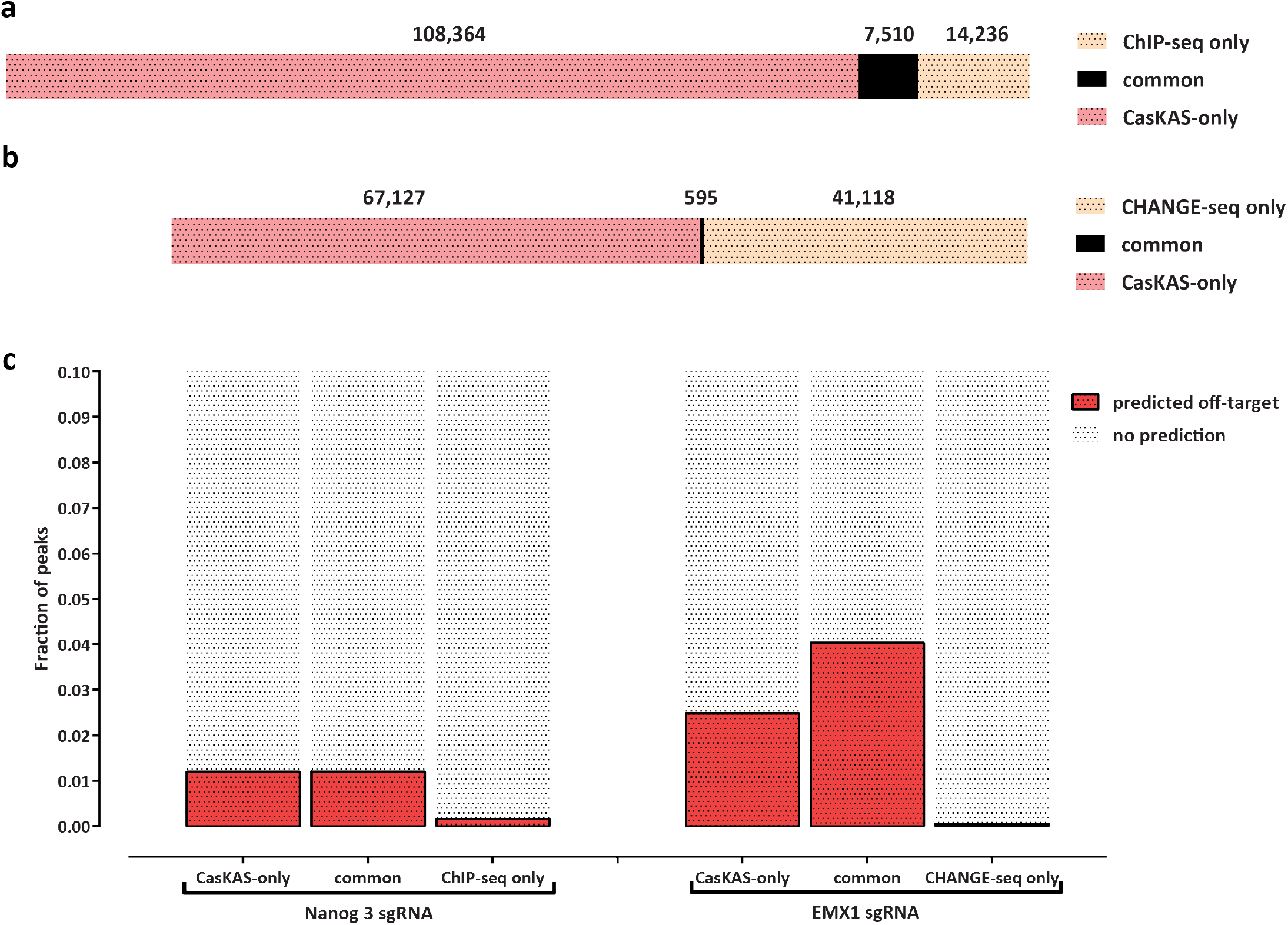
Comparing *in vitro* dCas9 results to using ChIP-seq and CHANGE-seq for off-target profiling. Shown is the overlap between MACS2 peak calls for the Nanog-sg3 sgRNA with Nanog ChIP-seq dataset (SRR1168384 from GEO accession ID GSE54745) in (a) and the EMX1 sgRNA with EMX1 CHANGE-seq (SRA accession SRX8227890) in (b). The fraction of peaks common or unique to each assay that are predicted to be off-targets for each sgRNA by Cas-OFFinder is shown in (c).

**Supplementary Figure 17:**
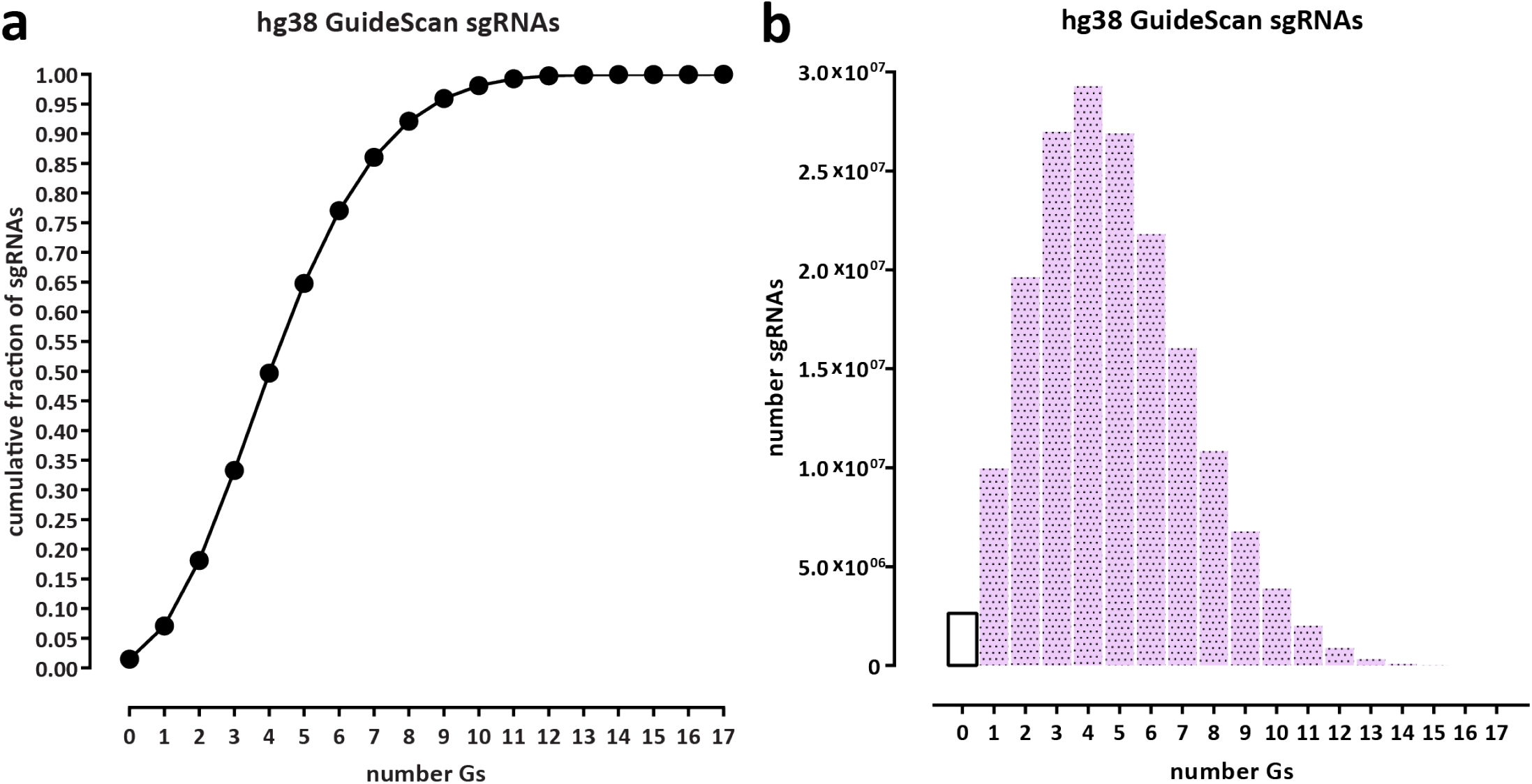
Most sgRNAs in the human genome contain multiple G nucleotides and are thus subject to labeling by N_3_-kethoxal. Statistics were calculated for all valid sgRNAs as defined by GuideScan ^18^ (a) Cumulative fraction of sgRNAs. (b) Absolute number of sgRNAs.

**Supplementary Figure 18:**
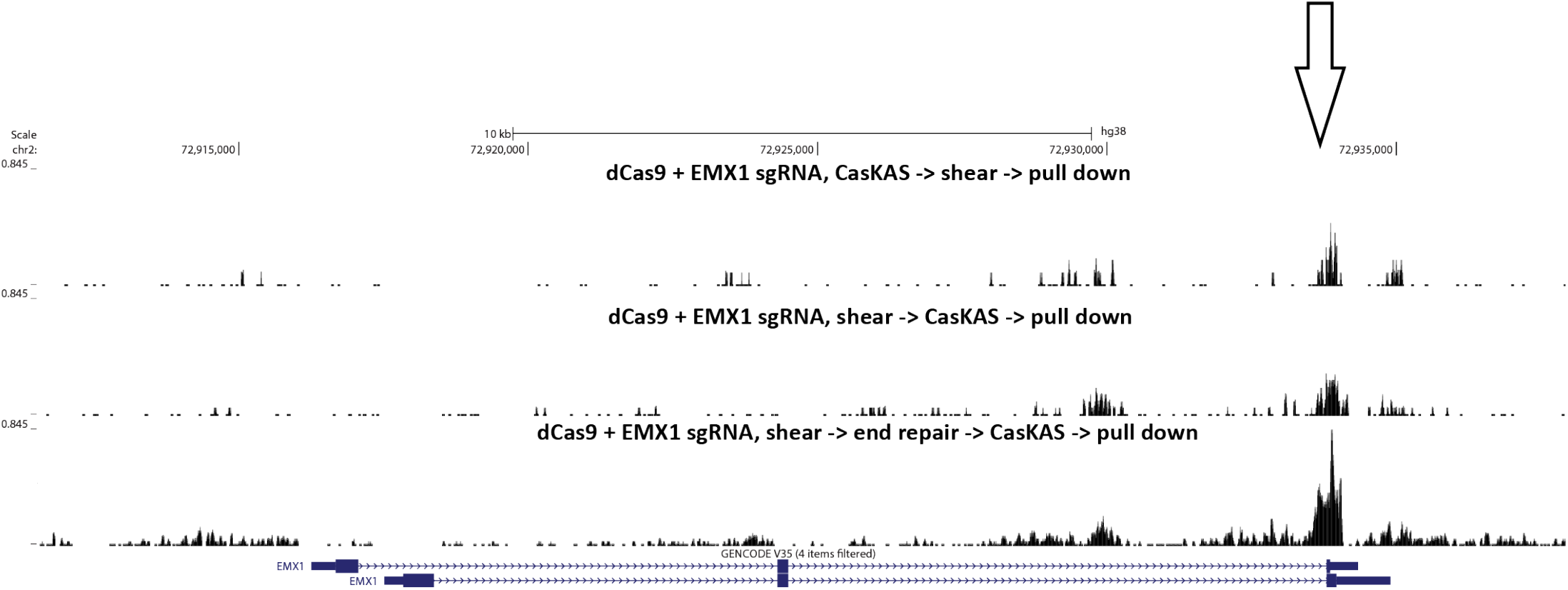
CasKAS can be performed on pre-sheared DNA. CasKAS was performed *in vitro* using the EMX1 sgRNA, first, conventionally, by carrying out the CasKAS reaction, then isolating and shearing genomic DNA, and also by pre-shearing the DNA and carrying out the CasKAS reaction on the fragmented DNA. The concern in that case is that the presence of sticky ends containing Gs and unprotected from the action of the N_3_-kethoxal would lower the background. This problem can be addressed by carrying out end repair on the sheared DNA prior to the CasKAS reaction.

